# GatorSC: Multi-Scale Cell and Gene Graphs with Mixture-of-Experts Fusion for Single-Cell Transcriptomics

**DOI:** 10.64898/2025.12.03.691688

**Authors:** Yuxi Liu, Zhenhao Zhang, Mufan Qiu, Song Wang, Flora D. Salim, Jun Shen, Tianlong Chen, Imran Razzak, Fuyi Li, Jiang Bian

## Abstract

Single-cell RNA sequencing (scRNA-seq) enables high-resolution characterization of cellular heterogeneity, but its rich, complementary structure across cells and genes remains underexploited, especially in the presence of technical noise and sparsity. Effectively leveraging this multi-scale structure is essentially an information fusion problem that requires integrating heterogeneous graph-based views of cells and genes into robust low-dimensional representations. In this paper, we introduce GatorSC, a unified representation learning framework that models scRNA-seq data through multi-scale cell and gene graphs and fuses them with a Mixture-of-Experts architecture. GatorSC constructs a global cell–cell graph, a global gene–gene graph, and a local gene–gene graph derived from neighborhood-specific subgraphs, and learns graph neural network embeddings that are adaptively fused by a gating network. To learn noise-robust and structure-preserving embeddings without labels, we couple graph reconstruction and graph contrastive learning in a unified self-supervised objective applied to both cell- and gene-level graphs. We evaluate GatorSC on 19 publicly available scRNA-seq datasets covering diverse tissues, species, and sequencing platforms. Experiments showed that GatorSC consistently outperforms state-of-the-art deep generative, graph-based, and contrastive methods for cell clustering, gene expression imputation, and cell-type annotation. The learned embeddings are used for accurate trajectory inference, recovery of canonical marker gene programs, and cell-type-specific pathway signatures in an Alzheimer’s disease single-nucleus dataset. GatorSC provides a flexible foundation for comprehensive single-cell transcriptomic analysis and can be readily extended to multi-omic and spatial modalities.

## Introduction

Single-cell RNA sequencing (scRNA-seq) has become a key technology for profiling transcriptomes at cellular resolution and for dissecting heterogeneous tissues, developmental processes, and disease-associated cell states [1, 2, 3]. However, scRNA-seq data are notoriously challenging to analyze: expression matrices are high-dimensional, extremely sparse due to pervasive dropout, and subject to substantial technical noise. scRNA-seq data inherently contains complementary structure at multiple scales, including global relationships between cells, functional dependencies among genes, and context-specific gene programs that are active only in particular cellular neighborhoods. Effectively exploiting this rich multi-scale structure is fundamentally an information fusion problem how to combine heterogeneous graph views into a single, robust representation of each cell.

A large body of work has applied deep representation learning to scRNA-seq data. Deep generative and autoencoder-based models such as scVI [4], scDeepCluster [5], DESC [6], and DeepBID [7] model count distributions with variational or deterministic autoencoders and couple reconstruction losses with clustering objectives, providing powerful latent representations for normalization, clustering, and denoising. Graph-based methods including scGNN [8], graph-sc [9], and scDSC [10] explicitly construct cell or gene graphs and utilize graph neural networks (GNNs) to propagate information across local neighborhoods, often achieving more accurate clustering and imputation than purely feature-based models. More recently, contrastive learning frameworks such as scGPCL [11] and scSimGCL [12] employ graph augmentations and instance- or prototype-level contrastive losses to learn noise-robust embeddings that are less tied to raw reconstruction errors. These approaches have substantially improved single-cell analysis, but they typically focus on one or two graph views and adopt relatively simple strategies for combining them.

Despite this progress, several important gaps remain from an information fusion perspective. First, most existing methods do not jointly model multi-scale structure across both cells and genes within a single representation. They may use a global cell–cell graph or a gene–cell bipartite graph, but rarely integrate a global cell manifold, a global gene network, and local, context-dependent gene dependencies. This makes it difficult to associate specific gene programs with particular regions of the cell manifold or to resolve closely related cell subtypes that differ by subtle shifts in gene expression programs. Second, when multiple graph views are considered, they are usually combined by fixed fusion schemes such as concatenation or uniform averaging, which cannot adapt the relative contribution of each view across datasets or even across individual cells. Third, most self-supervised objectives emphasize either reconstruction of noisy counts or contrastive agreement between graph views, but not both, potentially limiting the ability to simultaneously preserve graph topology, denoise expression, and enforce semantic consistency in the latent space.

In this study, we introduce **GatorSC**, a unified representation learning framework designed to fill these gaps by combining multi-scale graph modeling with a Mixture-of-Experts (MoE) fusion architecture and dual self-supervision. **GatorSC** constructs three hierarchical graphs from scRNA-seq data, including a global cell–cell graph capturing population-level organization, a global gene–gene graph encoding cross-cell functional dependencies, and a local gene–gene graph derived from neighborhood-specific subgraphs that emphasize context-dependent interactions, and then learns graph neural network embeddings on each graph. A modality-specific MoE assigns one expert to each graph view and uses a gating network to adaptively fuse expert outputs into a unified cell representation, allowing different datasets and cell populations to draw more heavily on the most informative structural signals. Finally, a unified self-supervised objective couples graph reconstruction losses with graph contrastive learning across the hierarchical graphs, jointly preserving structural fidelity and semantic consistency without relying on cell-type labels. Through extensive experiments on diverse scRNA-seq datasets, we empirically demonstrate that **GatorSC** yields robust gains in cell clustering, gene expression imputation, and cell-type annotation compared with state-of-the-art deep generative, graph-based, and contrastive methods, thereby establishing it as a flexible foundation for single-cell transcriptomic analysis.

## Methods

### Hierarchical Graph Modeling for scRNA-seq Data

In this paper, we propose a hierarchical graph modeling framework that progressively captures biological structure in single-cell data, ranging from local gene-level interactions within individual cells to global cellular population architectures, as illustrated in Figure 1A. Each layer builds upon the previous one while offering complementary biological insights. Accordingly, our framework consists of three hierarchical graphs that encode distinct yet synergistic levels of biological organization: a global cell–cell graph, a global gene–gene graph, and a local gene–gene graph.

**Figure 1.**
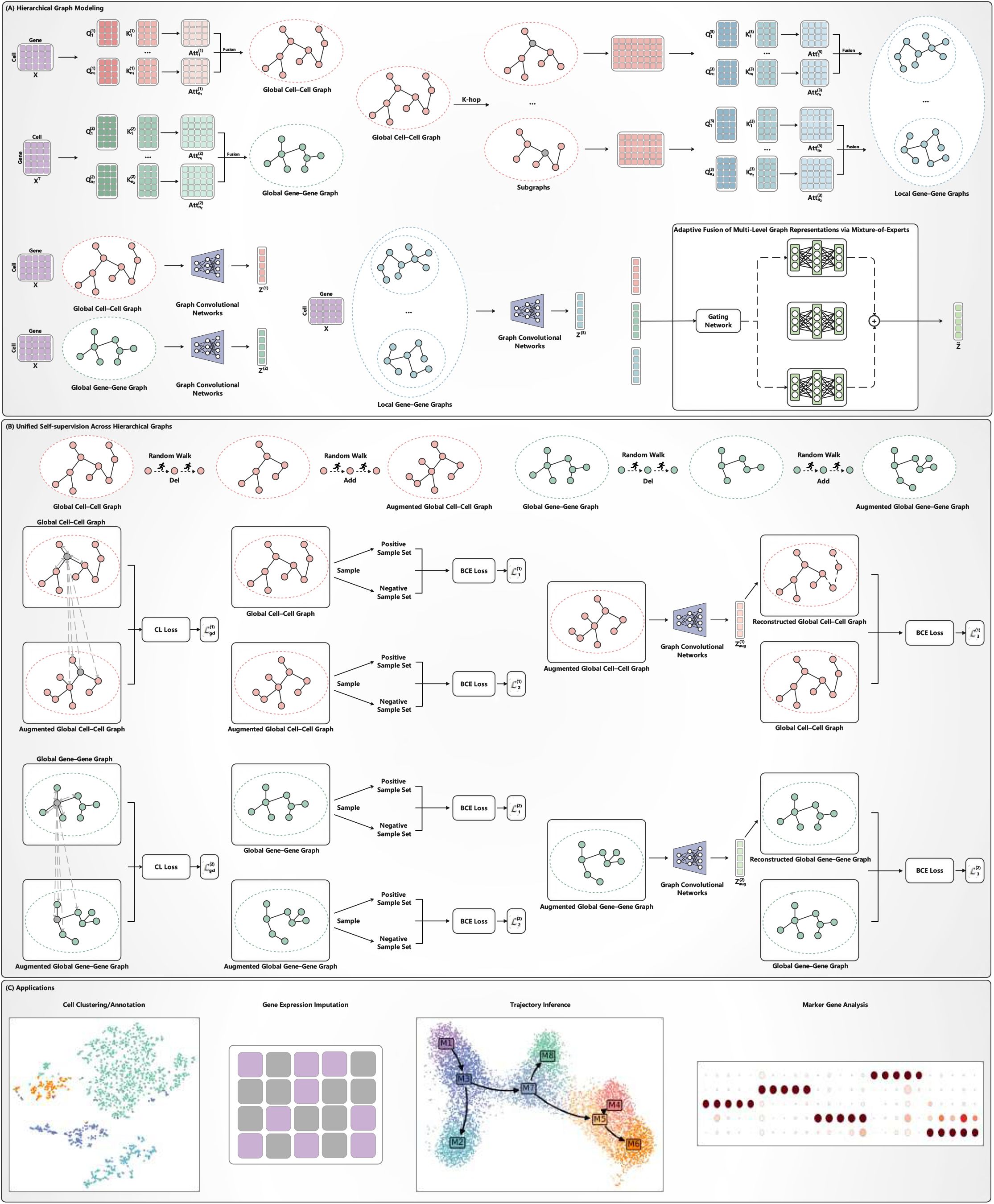
**Overview of the GatorSC framework**.

#### Global Cell–Cell Graph (Population-Level Structure)

Starting from the gene-expression matrix, we compute pairwise similarities between cells to construct a global cell–cell graph, which delineates the population-level architecture of the dataset. This graph captures global similarity patterns and large-scale cellular organization, serving as the structural backbone for downstream representation learning.

#### Global Gene–Gene Graph (Cross-Cell Dependency Network)

To capture gene-level dependencies that span multiple cells, we construct a global gene–gene graph that models cross-cell functional relationships among genes. This graph encodes robust gene–gene associations that remain stable across diverse cellular contexts. The resulting global gene network complements the cell–cell graph by providing a broader functional perspective, thereby bridging gene-level and cell-level representations.

#### Local Gene–Gene Graph (Context-Specific Dependency)

Building upon the global cell graph, we extract multiple cell subgraphs, each representing a localized cellular neighborhood. Within each subgraph, we recompute gene–gene similarities to construct context-specific gene graphs that captures dependency patterns unique to specific cellular contexts. These subgraph-level gene graphs are then integrated to form a local gene–gene graph that emphasizes fine-grained, context-dependent associations among genes, complementing the global graph structure with local specificity.

By integrating these three hierarchical levels of graphs (global cell–cell, global gene–gene, and local gene–gene), our framework offers a multi-scale, multi-view representation of scRNA-seq data. This design unifies both global and local dependencies across cells and genes, providing richer structural priors for deep learning models. Consequently, the learned representations are more biologically meaningful and better suited for downstream analyses, including cell clustering, cell type annotation, trajectory inference, and gene expression denoising.

##### Global Cell–Cell Graph

To formalize the above design, we introduce a unified notation for the three graphs. Let **X** ∈ ℝ^*M* ×*N*^ denote the gene expression matrix, where each row corresponds to a cell and each column to a gene, with *M* being the number of cells and *N* the number of genes. From **X**, we construct three graphs: the global cell–cell graph 𝒢^(1)^, the global gene–gene graph 𝒢^(2)^, and the local gene–gene graph 𝒢^(3)^. For *i* = 1, 2, 3, we write

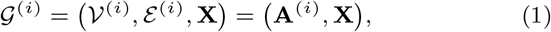

where 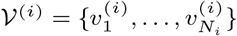 is the node set, ℰ ^(*i*)^ is the edge set, and 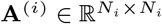 is the corresponding adjacency matrix.

To construct the global cell–cell graph 𝒢^(1)^, we compute its adjacency matrix **A**^(1)^ using a multi-head attention mechanism to learn cell–cell similarities. Given the gene expression matrix **X** ∈ ℝ^*M* ×*N*^, we perform linear projections to obtain the query and key for each attention head. Then, scaled dot-product attention is applied to compute pairwise similarities between cells, followed by Softmax normalization to obtain attention weights. Each attention head is computed independently, and the resulting outputs are concatenated and linearly transformed to obtain the adjacency matrix. The mathematical formulation can be written as follows:

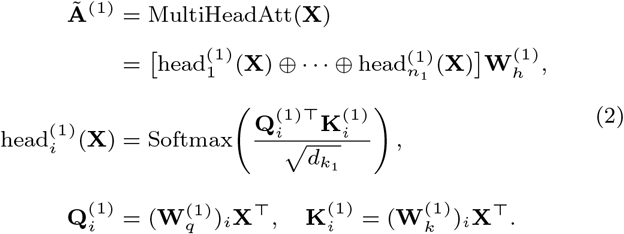

where all **W** is trainable parameters. 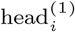 is the i-th attention head. *n*_1_ is the number of heads. ⊕ is the concatenation. 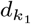 is the dimension of **K**^(1)^. Accordingly, a sparse and non-negative adjacency matrix **A**^(1)^ can be obtained using the intermediate matrix **Ã** ^(1)^ with numerical constraints as follows:

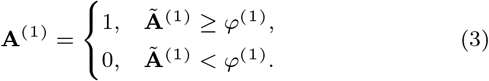

’
where *φ*^(1)^ is a learnable threshold that excludes the values lower than that.

##### Global Gene–Gene Graph

We construct a global gene–gene graph 𝒢^(2)^ by computing its adjacency matrix **A**^(2)^ at the gene level. To learn the intrinsic relationships among genes, we use the transposed gene expression matrix **X**^⊤^ ∈ ℝ^*N* ×*M*^, and apply a multi-head scaled dot-product attention mechanism, consistent with that used in the previous section, to compute pairwise similarity weights among genes. The outputs from all attention heads are aggregated to obtain the adjacency matrix. The mathematical formulation can be written as follows:

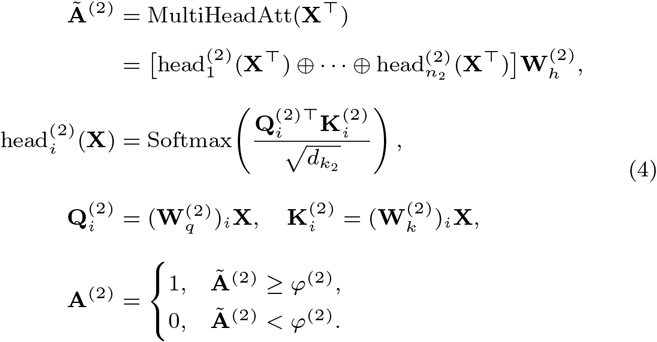

where all **W** is trainable parameters. *n*_2_ is the number of attention heads. 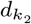 is the dimension of **K**^(2)^. *φ*^(2)^ is a learnable threshold that excludes the values lower than that.

##### Local Gene–Gene Graph

Based on the foundation established by the global cell–cell graph, a subgraph is extracted for each node to represent its structural relationships. In particular, the subgraph for a given node consists of its k-hop neighboring nodes. We take an example for node 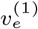, and its subgraph 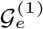 can be defined as follows:

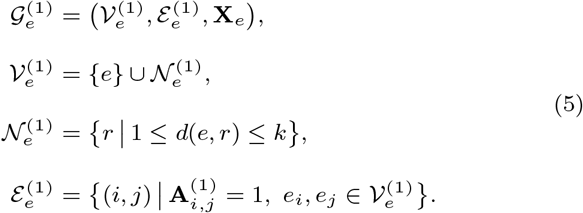

where 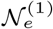 is the set of k-hop neighboring nodes of 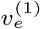. Through the above steps, given a node 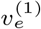, we can obtain its subgraph 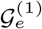 and the corresponding set of k-hop neighbors 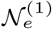. Based on 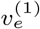 and 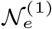, we can extract the subset of gene expression data corresponding to node 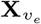. Using 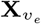 as input, we then apply the multi-head scaled dot-product attention mechanism again, to compute the adjacent matrix. The mathematical formulation can be written as follows:

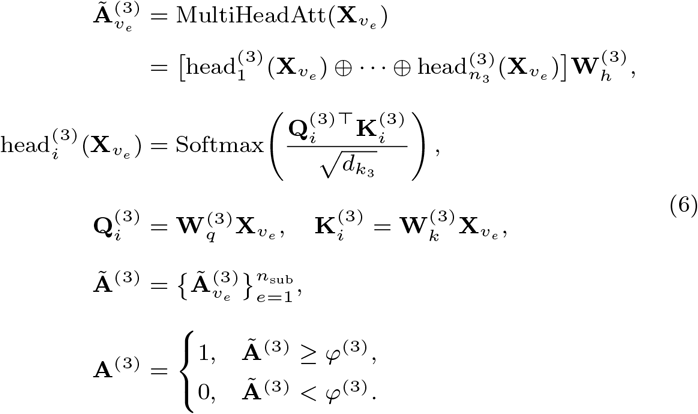

where all **W** is trainable parameters. *n*_3_ is the number of attention heads. 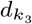 is the dimension of **K**^(3)^. *n*_sub_ is the number of subgraphs. *φ*^(3)^ is a learnable threshold that excludes the values lower than that.

##### Adaptive Fusion of Multi-Level Graph Representations via Mixture-of-Experts

After obtaining the three graphs 𝒢^(1)^, 𝒢^(2)^, 𝒢^(3)^, the next step is to learn the node representations 𝒱^(*i*)^ in each graph, using Graph Convolutional Networks (GCNs) and Multilayer Perceptron (MLP), as formulated below:

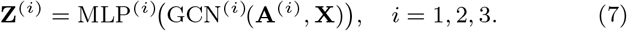

where **Z**^(*i*)^ is the learned representation of **X** and 𝒢^(*i*)^. The obtained representations **Z**^(1)^, **Z**^(2)^, **Z**^(3)^ capture three distinct levels of intrinsic data dependencies. These representations are complementary rather than redundant. Since each representation emphasizes different structural aspects, a naive fusion strategy such as simple concatenation or averaging would ignore the varying relative importance of these levels. We introduce an adaptive fusion mechanism based on the MoE paradigm. MoE is a modular neural architecture in which multiple specialized experts are trained to model different input contributions, while a gating network adaptively controls their weight [13]. Instead of relying on a single network to handle all variations in the data, MoE introduces a gating network that dynamically assigns input samples to different experts based on their characteristics [14]. The gating network produces a probability distribution over experts, enabling the model to adaptively weigh the contributions of each expert and combine heterogeneous representations in a flexible and data-driven manner. Formally, we employ an MoE framework that consists of a set of *L* expert networks (with *L* = 3 in this study) and a gating network. Let *G*(**X**) denote the output of the gating network and *f*_*i*_(**X**) the output of the i-th expert network for a given input **X**. The aggregated dense representation generated by the MoE can be written as:

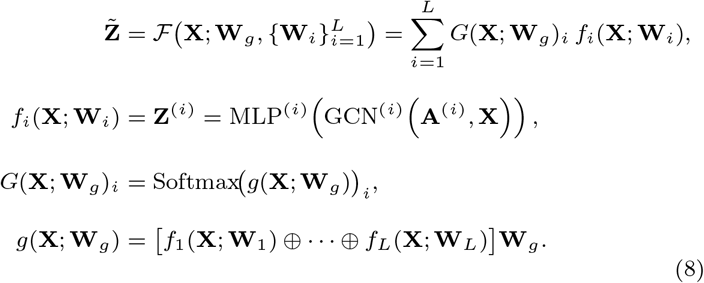

where 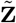 is the dense representation learned. All **W** are trainable parameters. *L* is the number of expert networks, which is set to *L* = 3.

### Unified Self-supervision Across Hierarchical Graphs

After obtaining the learned dense representation, we further optimize it in an end-to-end manner through a unified self-supervised learning framework that integrates both *generative* and *contrastive* paradigms, as shown in Figure 1B. Self-supervised learning approaches can be broadly categorized into two types. The generative objective focuses on reconstructing the original signal or graph structure (e.g., graph reconstruction), emphasizing structural fidelity [15]. By recovering missing or corrupted connections, this approach enhances the model’s robustness to incomplete or noisy data and benefits downstream tasks such as denoising, imputation, and link prediction. In contrast, the contrastive objective encourages consistent representation among semantically identical instances under different perturbations or graph views, while maintaining uniformity in the embedding space [16]. This helps the model learn noise-invariant and discriminative representations, improving performance on clustering, classification, and related tasks.

We design two self-supervised objectives: graph reconstruction learning and graph contrastive learning. Before defining these objectives, we introduce graph augmentation, a critical step that generates semantically consistent yet structurally diverse graph views. Graph augmentation serves two primary purposes: (i) providing controlled corrupted inputs for generative tasks (e.g., node masking or edge removal), to enhance structural learning, and (ii) creating positive and negative sample pairs for contrastive learning to promote invariance and prevent representation collapse under perturbations.

We modify the graph structure by performing path-level operations, including edge deletion and addition along the sampled paths, to generate structurally perturbed views of the original graph. Given the graph 𝒢^(1)^, the augmentation process can be defined as follows:

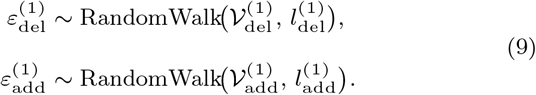

where 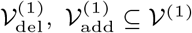 are sets of root nodes sampled from graph 𝒢^(1)^ follow Bernoulli distributions, i.e., 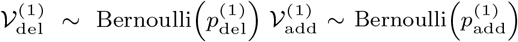, where 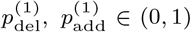 1) are the dele and addition probabilities, and 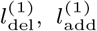are the maximum path lengths. Each set 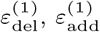 consists of edges/paths encountered by random walks starting from the sampled root nodes. A path is a sequence of edges; therefore, 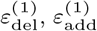 can be viewed as set of edges. Based on these sets, we construct the augmented graph as follows:

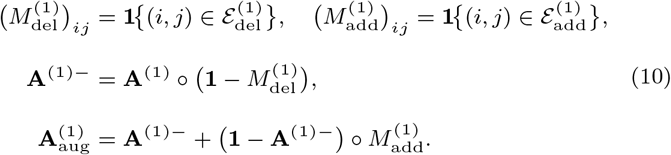

where the symbol º denotes the Hadamard (element-wise) product; **1** is the all-ones matrix of the same size as **A**^(1)^; all additions/subtractions are element-wise. 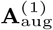 is the adjacent matrix of the augmented graph, which we denote by 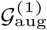. Having obtained the original graph 𝒢^(1)^ and its augmented version 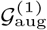,we can now define two self-supervised tasks: graph reconstruction and graph contrastive learning.

We first introduce the graph reconstruction task. Specifically, the set of observed edges in the original graph 𝒢^(1)^ is denoted as ℰ^(1)+^, which serves as the positive sample set, representing pairs of nodes that should be placed closer together in the embedding space. Conversely, a set of randomly sampled unobserved node pairs from 𝒢^(1)^ is used as the negative sample set, denoted as ℰ ^(1)−^. Similarly, for the augmented graph 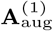, we define the corresponding positive and negative edge sets as 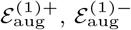, respectively. Based on these positive and negative edge sets, the reconstruction loss can be formulated as follows:

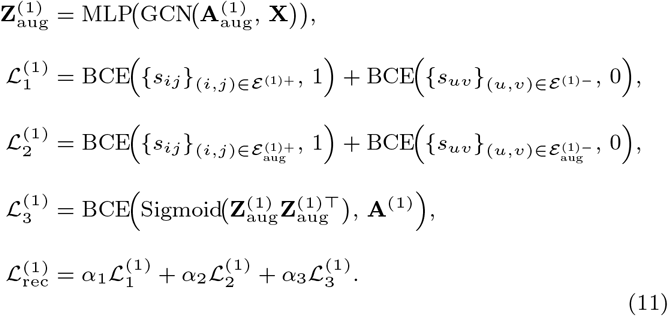

where *s*_*ij*_ = **x**_*i*_ · **x**_*j*_ is the dot-product 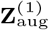 is the learned representation of **X** in the augmented graph 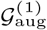. BCE(·) represents the binary loss of cross-entropy.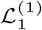 and 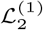 correspond to the reconstruction losses for 𝒢^(1)^ and 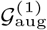, respectively. These loss terms encourage the model to assign higher similarity scores to observed (positive) edges while assigning lower scores to randomly sampled non-edges. 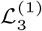 denotes the loss in reconstructing 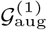 from 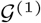. *α*_1_, *α*_2_, *α*_3_ are coefficients that balance the contributions of the three losses. The total reconstruction loss is defined as 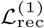.

Next, we introduce the graph contrastive learning task. The core idea of contrastive learning is to define positive and negative sample pairs and optimize the model to maximize the similarity of positive pairs while minimizing the similarity of negative pairs. Based on 𝒢^(1)^ and its augmented counterpart 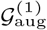, we define the positive samples as follows: a node from 𝒢^(1)^ is selected as an anchor. The corresponding positive samples include: (i) the nodes directly connected to the anchor in 𝒢^(1)^, (ii) its counterpart node in 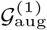, and (iii) the nodes connected to the anchor from 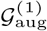, while all remaining nodes are treated as negative samples. Given these definitions, the contrastive loss can be formulated as:

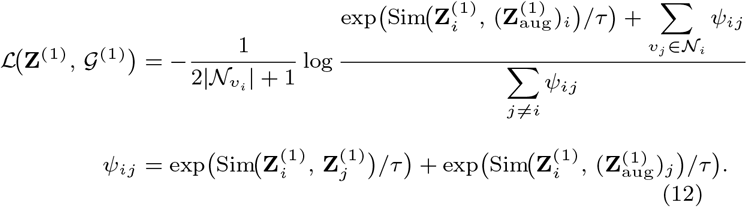

where sim(·) denotes the similarity function, implemented as a dot product, and *τ* is a temperature parameter that controls the strength of the penalty for negative pairs. 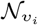 is a set of positive neighbors for the i-th node in ℒ(**Z**^(1)^, 𝒢^(1)^) represents the contrastive loss when the anchor is selected from 𝒢^(1)^. Similarly, when the anchor is selected from 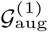, the contrastive loss is denfoted as 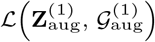. The overall contrastive loss 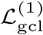 is then defined as:

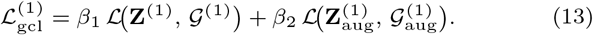

where *β*_1_, *β*_2_ are coefficients that balance the contributions of the two contrastive losses.

Based on the above steps, the overall self-supervised loss for each hierarchical graph 𝒢^(1)^ can be defined as:

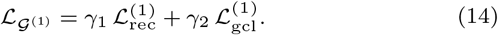

where *γ*_1_, *γ*_2_ are coefficients that balance the generative (reconstruction) and contrastive objectives.

In summary, 𝒢^(1)^ models cell–cell relationships from a global population perspective, while 𝒢^(2)^ captures gene–gene dependencies from a global gene-level viewpoint. Based on 𝒢^(1)^, we can obtain the corresponding self-supervised loss 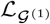; and based on 𝒢^(2)^, we can obtain 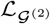. Finally, the overall self-supervised loss that integrates both cell and gene perspectives is formulated as:

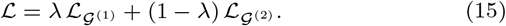

where *λ* is a trade-off coefficient that controls the relative importance of the two components. This unified objective allows the model to simultaneously preserve global structural fidelity (through reconstruction) and semantic consistency (through contrastive learning) across both cell- and gene-level graphs.

## Application tasks

As described above, the model is trained by minimizing the loss function defined in Eq. (15). After training, the matrix 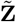 in Eq. (8) serves as the learned low-dimensional representation of the cells. Building on this unified representation, our framework can be readily applied to a variety of downstream single-cell analysis tasks, as illustrated in Figure 1C.

For the clustering task, we feed 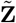 into a general clustering method to obtain cluster assignments, where we use k-means as our clustering algorithm.

For the imputation task, we use 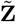 as input to a regressor. Specifically, we employ a fully connected layer as the regressor to predict and impute missing gene expression values as:

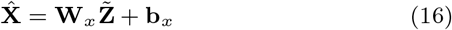

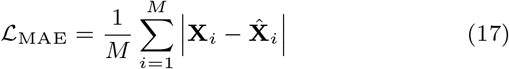

where **W**_*x*_ and **W**_*x*_ are trainable parameters. 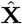 denotes the imputed gene expression matrix. The mean absolute error was used as the loss function to optimize the imputation task.

For the cell-type annotation task, we use 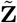 as input to a multi-class classifier. Specifically, we employ a fully connected layer to predict cell-type labels as:

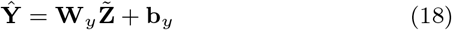

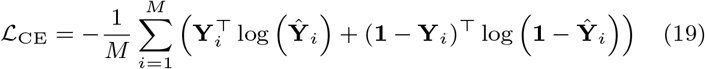

where **W**_*y*_ and **b**_*y*_ are trainable parameters, and **Ŷ** denotes the predicted cell-type labels. The cross-entropy loss was used as the loss function to optimize the cell-type annotation task.

## Results

### Benchmark datasets and Baselines

We evaluated **GatorSC** on three key single-cell analysis tasks, including cell clustering, gene expression imputation, and automated cell-type annotation, using multiple publicly available scRNA-seq datasets spanning diverse tissues, species, and experimental platforms [17, 18, 19, 20, 21, 22, 23, 24, 1, 25], as shown in Table 1. **Details on data preprocessing, implementation and parameter settings, evaluation metrics, visualization results, and additional experimental results, as well as ablation studies and hyperparameter sensitivity analyses, are provided in the Supplementary Materials**.

**Table 1.** Summary of scRNA-seq benchmark datasets, sequencing platforms, and basic statistics.

| Dataset | Sequencing platform | # of Cells | # of Genes | # of cell types |
| --- | --- | --- | --- | --- |
| BCs-PCs [17] | 10x 5' v2 | 3912 | 19283 | 6 |
| Chen [18] | 10x 3' v3 | 36313 | 20043 | 15 |
| Emont [19] | 10x 3' v3 | 7632 | 24811 | 4 |
| Global [17] | 10x 5' v2 | 10831 | 19283 | 31 |
| HTAPP-213-SMP-6752 [20] | 10x 3' v3 | 9799 | 24855 | 7 |
| HTAPP-313-SMP-932 [20] | 10x 3' v3 | 12494 | 20938 | 10 |
| HTAPP-783-SMP-4081 [20] | 10x 3' v2 | 2276 | 16853 | 8 |
| Quake-Diaphragm [1] | Smart-seq2 | 870 | 23341 | 5 |
| Quake-Limb Muscle [1] | Smart-seq2 | 1090 | 23341 | 6 |
| Quake-Lung [1] | Smart-seq2 | 1676 | 23341 | 11 |
| Quake-Trachea [1] | Smart-seq2 | 1350 | 23341 | 4 |
| Zeisel [21] | STRT-seq UMI | 3005 | 19972 | 9 |
| Mouse ES [22] | inDrop | 2717 | 24175 | 4 |
| Pancreas [23] | inDrop | 1937 | 15575 | 14 |
| AMB3 [24] | SMART-Seq v4 | 12832 | 42625 | 3 |
| AMB16 [24] | SMART-Seq v4 | 12832 | 42625 | 16 |
| AMB92 [24] | SMART-Seq v4 | 12832 | 42625 | 92 |
| TM [1] | SMART-Seq2 | 54865 | 19791 | 55 |
| Zheng [25] | 10x Chromium | 65943 | 20387 | 11 |

Across clustering, imputation, and annotation, we benchmarked **GatorSC** against representative deep generative/autoencoder models (DESC [6], scDeepCluster [5], scVI [4], DeepBID [7], DCA [26], AutoClass [27]), graph-based methods (scGNN [8], graph-sc [9], scDSC [10], GE-Impute [28], MAGIC [29], scBFP [30]) and graph contrastive learning methods (scGPCL [11], scSimGCL [12]), classical clustering algorithms (KMeans, Leiden [31], Louvain [32]), and supervised/atlas-based annotators (scHeteroNet [33], scDeepSort [34], ACTINN [35], CellTypist [36], scANVI [37]).

### Cell clustering performance on 14 benchmark scRNA-seq datasets

We evaluated **GatorSC** on the cell clustering task using 14 benchmark scRNA-seq datasets spanning diverse tissues, species, and sequencing platforms. As summarized in Figure 2, **GatorSC** consistently outperforms representative deep generative, graph-based, and conventional clustering methods across these datasets. Across four metrics, adjusted Rand index (ARI), normalized mutual information (NMI), clustering accuracy (ACC), and homogeneity, **GatorSC** achieves the best or second-best scores the best or second-best scores. The advantage is especially notable on heterogeneous datasets with many cell types, such as Global (31 cell types) and Pancreas (14 cell types). Graph-based baselines, such as scGPCL and scSimGCL, often outperform purely autoencoder-based approaches (DESC, scDeepCluster, DeepBID), highlighting the importance of explicitly modeling cell–cell structure. Nevertheless, no single baseline dominates across all datasets or metrics, whereas **GatorSC** remains among the top-performing methods in most cases. In contrast, classical clustering algorithms (KMeans, Leiden, and Louvain) generally underperform and show larger performance fluctuations. Overall, these results suggest that the hierarchical graph modeling and Mixture-of-Experts fusion in **GatorSC** yield more discriminative and biologically meaningful representations by integrating global cell–cell structure, global gene–gene relationships, and local context-specific gene–gene interactions, leading to robust performance across diverse settings.

**Figure 2.**
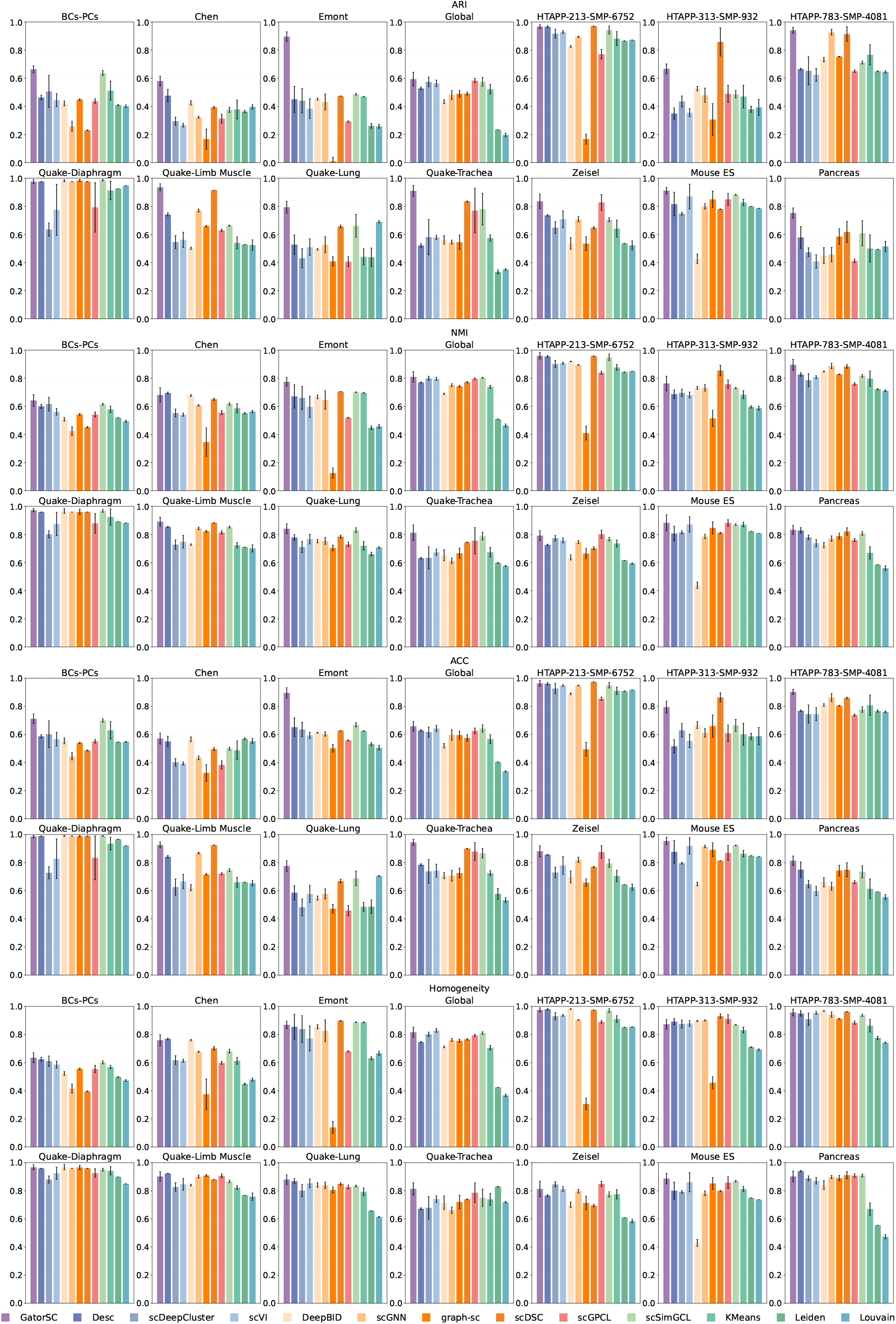
**Clustering performance of GatorSC and baseline methods on 14 benchmark scRNA-seq datasets. For each dataset, rows show ARI, NMI, ACC, and homogeneity scores for GatorSC and baseline methods; higher values indicate better clustering performance**.

### Gene expression imputation performance on 14 benchmark scRNA-seq datasets

We evaluated the gene expression imputation performance of **GatorSC** on the same 14 benchmark scRNA-seq datasets under simulated dropout rates from 10% to 50%. As shown in Figure 3, **GatorSC** consistently achieves higher Pearson correlation coefficients (PCC) and lower L1 errors between imputed and ground-truth expression than competing denoising and imputation methods across most datasets and dropout levels. Improvements are observed in relatively simple datasets with few cell types (e.g., Emont, Mouse ES) and highly heterogeneous datasets such as Global and Pancreas, indicating that **GatorSC** can recover expression signals under complex population structure. Notably, the advantage becomes more pronounced as dropout increases: at mild dropout (10–20%), scSimGCL is competitive on several datasets, whereas under severe dropout (40–50%) many baselines degrade substantially, particularly in PCC, while **GatorSC** remains relatively stable across datasets. These results suggest that the hierarchical graph construction and Mixture-of-Experts fusion enable **GatorSC** to leverage both local and global transcriptomic structure to infer missing values, even when the expression matrix is highly sparse. Overall, **GatorSC** serves as a strong clustering model and an effective imputation method, learning representations that preserve global correlation patterns while accurately reconstructing local expression profiles.

**Figure 3.**
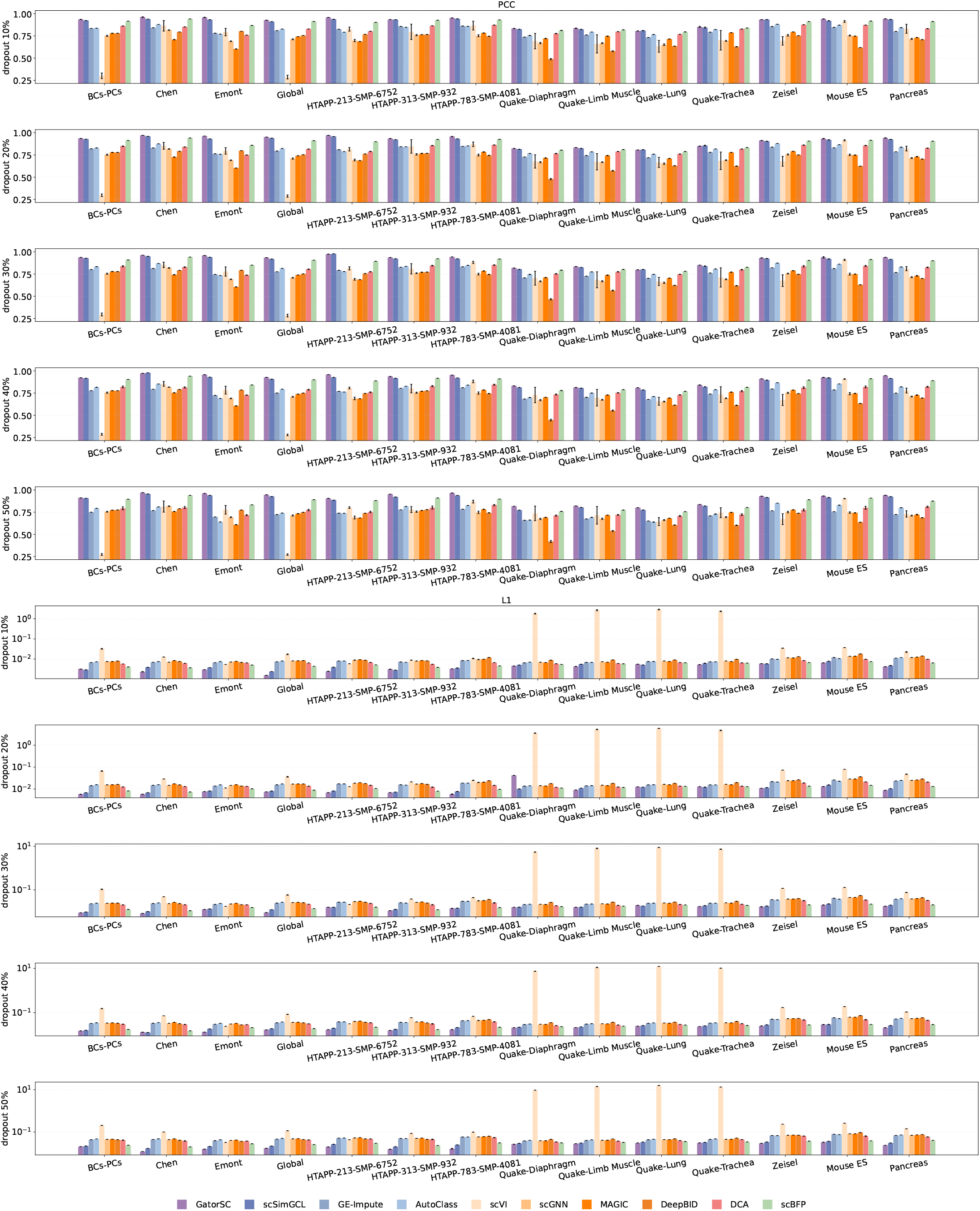
**Gene expression imputation performance of GatorSC and competing methods on 14 scRNA-seq datasets under increasing simulated dropout rates (10–50%). The top panels report Pearson correlation coefficients (PCC) between imputed and ground-truth expression, and the bottom panels report L1 errors. Higher PCC and lower L1 values indicate better imputation performance**.

### Cell-type annotation performance on 14 benchmark scRNA-seq datasets

We further evaluated **GatorSC** on the cell-type annotation task using 14 benchmark scRNA-seq datasets with ground-truth labels. As shown in Figure 4, **GatorSC** consistently matches or outperforms widely used supervised and semi-supervised classifiers, including scHeteroNet, ACTINN, scDeepSort, CellTypist, and scANVI. Across three metrics, F1 score, overall accuracy (ACC), and Matthews correlation coefficient (MCC), **GatorSC** achieves the best or near-best performance on most datasets, suggesting that the learned representations transfer well to downstream classification. Compared with strong graph-based baselines, scHeteroNet attains solid performance on several datasets, yet **GatorSC** provides consistent gains on many of them, particularly on heterogeneous datasets such as Global and Pancreas. On simpler datasets with only a few broad cell types (e.g., BCs-PCs and Emont), **GatorSC** performs comparably to the best baselines. These results indicate that the hierarchical graph modeling and Mixture-of-Experts fusion in **GatorSC** yield representations that are effective for both unsupervised clustering and supervised annotation. By jointly leveraging cell–cell and gene–gene structure in a unified self-supervised framework, **GatorSC** supports accurate and robust cell-type labeling across diverse single-cell transcriptomic studies.

**Figure 4.**
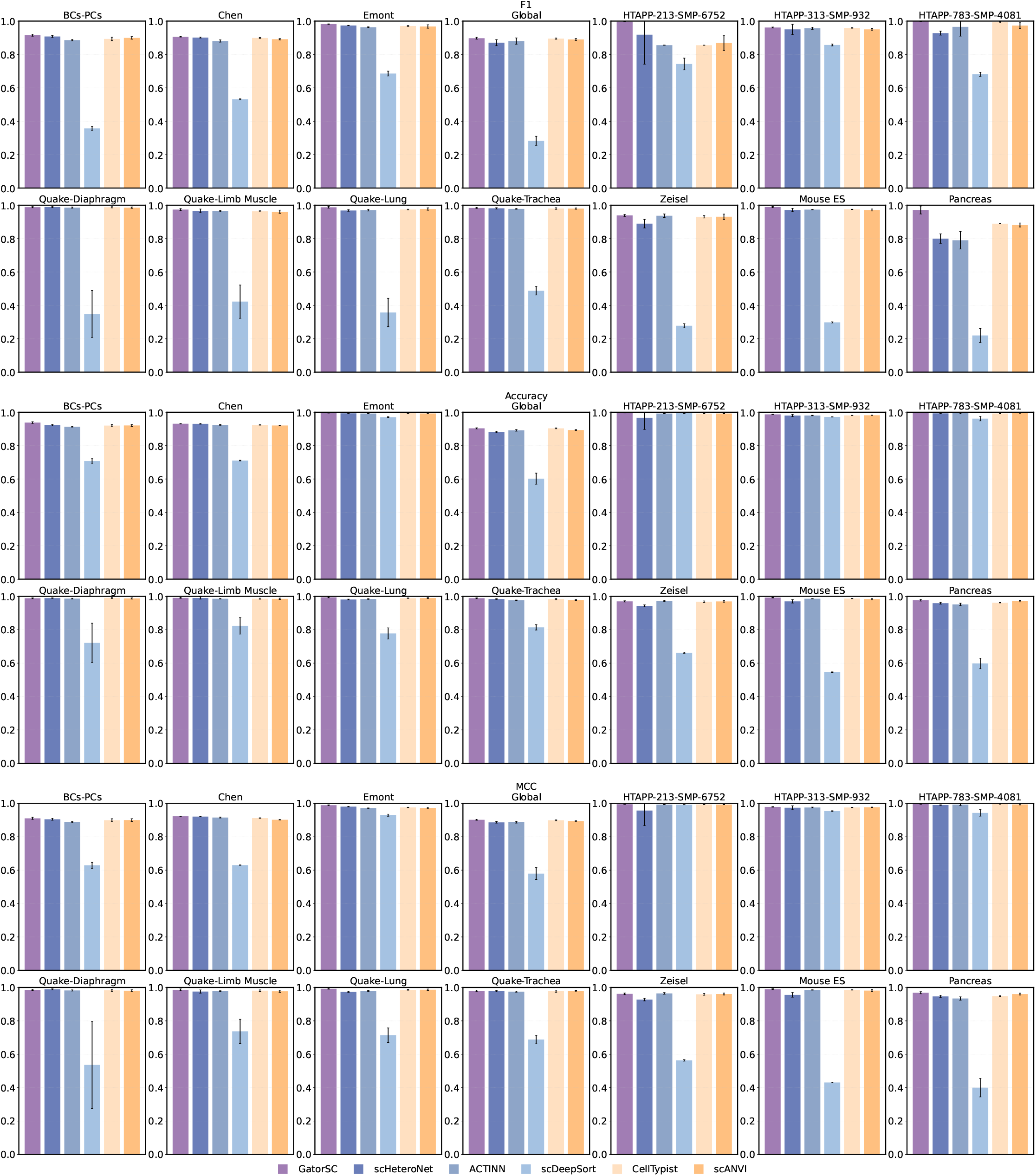
**Cell-type annotation performance of GatorSC and competing methods on 14 scRNA-seq datasets. Bars show F1 score (top row), overall accuracy (middle row), and Matthews correlation coefficient (MCC, bottom row) for GatorSC and competing methods. Higher values of all three metrics indicate better cell-type annotation performance**.

### Trajectory inference on the binary_tree_8 benchmark dataset

Figure 5 summarizes the developmental trajectory inferred by **GatorSC** from the latent representation of the `binary_tree_8` single-cell transcriptomes [38]. **GatorSC** first aggregates cells into eight meta-states (M1–M8) and then orders both meta-states and individual cells along a branching developmental continuum. In the Topology panel, the meta-states are arranged as a directed tree that captures the global branch structure of the trajectory. M1 is identified as an upstream progenitor-like state that flows into M3 and subsequently bifurcates into several branches. From M3, one branch terminates in M2, whereas another proceeds through M7 to M5, which further splits into the terminal states M4 and M6; a separate branch from M3 ends in M8. This topology is consistent with the underlying design of the `binary_tree_8` benchmark and highlights the main decision points and terminal fates inferred by **GatorSC**.

**Figure 5.**
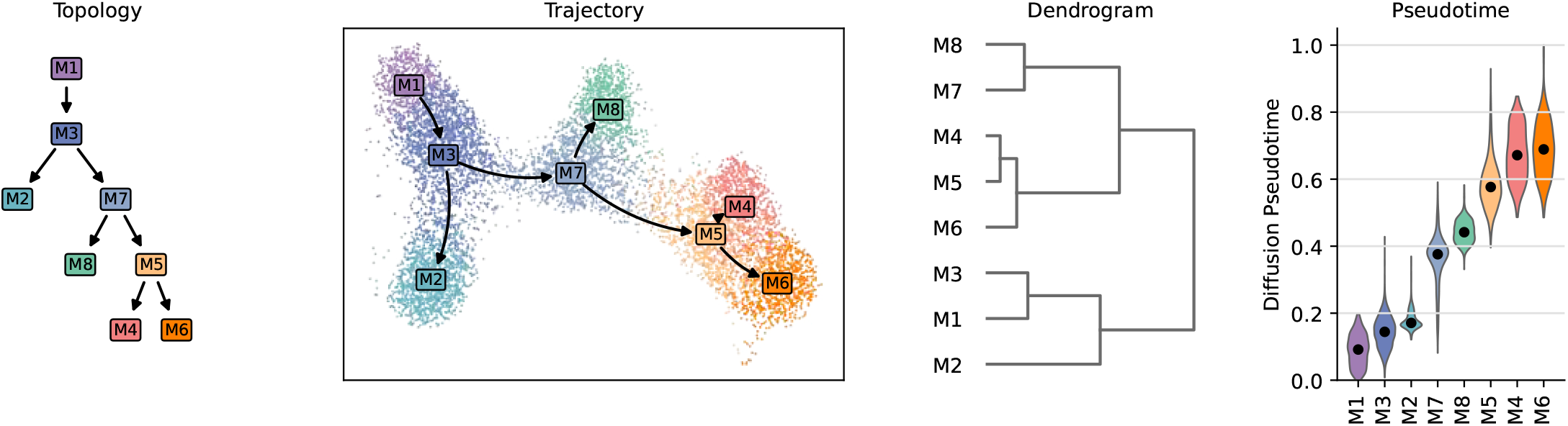
**Trajectory inference on the** `binary_tree_8` **benchmark dataset by GatorSC. The Topology panel (left) shows the directed tree connecting eight meta-states (M1–M8) and recapitulates the ground-truth branching structure. The Trajectory panel visualizes single cells in the learned low-dimensional manifold, colored by meta-state, with the black curve indicating the principal developmental paths from the upstream state M1 toward downstream branches. The Dendrogram panel depicts hierarchical clustering of meta-states based on their transcriptomic similarity, closely matching the inferred branch structure. The Pseudotime panel displays diffusion pseudotime distributions for each meta-state, revealing a gradual increase from early states (M1, M3, M2) to late states (M5, M4, M6), consistent with the inferred developmental progression**.

In the Trajectory panel, each point represents a single cell in the low-dimensional manifold and colors denote the inferred meta-states. The black curve traces the principal trajectory learned by **GatorSC**, with arrows indicating progression from M1 toward downstream branches. Cells form continuous clouds that smoothly connect neighboring meta-states along each branch, and branch junctions are populated by cells assigned to intermediate states such as M3 and M7. This spatial organization indicates that the discrete meta-state structure is coherent with an underlying continuous manifold of cell states rather than being driven by isolated clusters.

The Dendrogram panel depicts the hierarchical relationships among meta-states based on their transcriptomic similarity. Early states (M1, M3, and M2) cluster together and are clearly separated from the group comprising late states (M5, M4, and M6), whereas M7 and M8 form an intermediate branch. This clustering pattern closely mirrors the branching structure in the topology panel, supporting the view that the inferred branches correspond to genuine transcriptional diversification of cell populations.

The Pseudotime panel shows the distribution of diffusion pseudotime values for each meta-state. Early meta-states (M1, M3, and M2) exhibit low pseudotime values, intermediate states (M7 and M8) occupy central pseudotime ranges, and late states (M5, M4, and M6) concentrate at high pseudotime values. The monotonic increase in median pseudotime from M1 to downstream meta-states, with limited overlap between early and late distributions, suggests that **GatorSC** recovers a temporally consistent ordering of cell states that is aligned with the inferred trajectory topology.

These results demonstrate that **GatorSC** reconstructs a developmental continuum consistent with the known branching structure of the `binary_tree_8` dataset, in which an upstream progenitor-like state (M1) progresses through intermediate states and diverges into multiple terminal branches. The agreement between topology, low-dimensional embedding, hierarchical relationships, and pseudotime supports the robustness of the inferred trajectory.

### Marker gene analysis on the Quake Smart-seq2 diaphragm dataset

To evaluate whether **GatorSC** recovers biologically meaningful structure, we examined the expression of canonical marker genes across major cell populations in the Quake Smart-seq2 diaphragm dataset [1] (Figure 6). In the dot plot (Figure 6A), a curated panel of lineage-specific markers is shown for **GatorSC**-annotated skeletal muscle satellite stem cells, mesenchymal stem cells, endothelial cells, macrophages, and lymphocytes. Dot size encodes the fraction of cells in each group expressing a given gene, and dot color represents the mean expression level. Stromal and mesenchymal markers such as MFAP5, CLEC3B, COL1A2, LUM, and DCN are enriched in the mesenchymal stem cell population, whereas endothelial-associated genes including CD36, FABP4, GPIHBP1, and CDH5 show selective expression in the endothelial compartment. Myeloid markers such as FCER1G, CCL9, CTSC, CTSS, and RAC2 are concentrated in the macrophage cluster, while the lymphocyte marker CD52 is largely restricted to the lymphocyte population, indicating largely lineage-restricted expression patterns.

**Figure 6.**
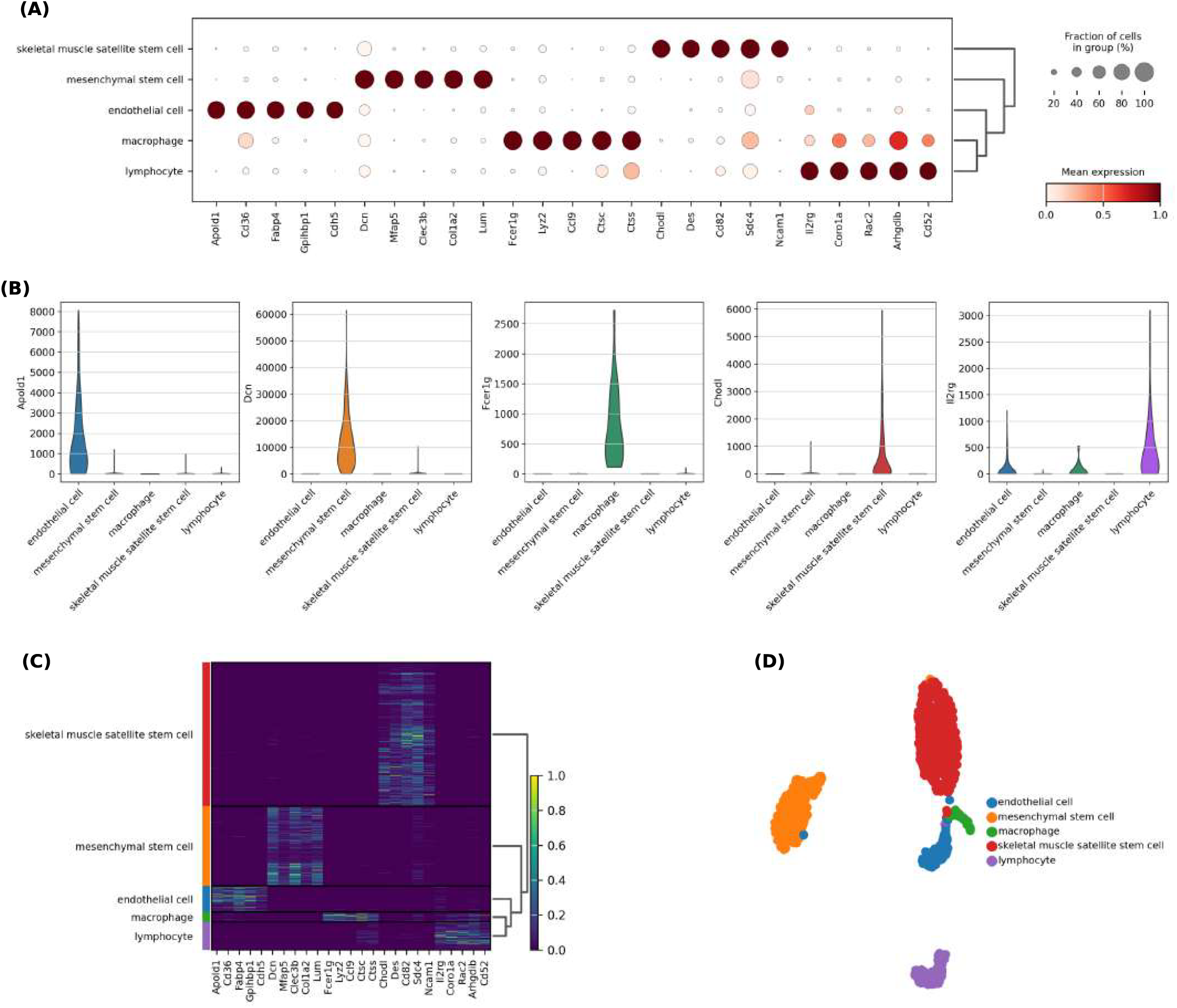
**Marker gene analysis in the Quake Smart-seq2 diaphragm dataset. (a) Dot plot of average expression of selected canonical marker genes across five GatorSC-annotated cell types. (b) Violin plots of representative marker gene expression in each cell type. (c) Heatmap of normalized marker scores summarizing the marker panel. (d) UMAP visualization of the GatorSC latent representation, colored by inferred cell-type labels**.

To provide complementary summaries of these markers, violin plots (Figure 6B) depict the distribution of expression for representative genes across the five **GatorSC**-defined cell types, and a heatmap of normalized marker scores (Figure 6C) aggregates signal across the full marker panel. In each case, the corresponding cell type shows higher expression and a larger fraction of marker-positive cells than unrelated populations, which show low signal. Finally, a two-dimensional embedding of the **GatorSC** representation (Figure 6D), with cells colored by inferred label, reveals well-separated clusters corresponding to the five major cell types. These results demonstrate that **GatorSC** yields coherent cell-type annotations that recapitulate known skeletal muscle and immune cell biology and support marker-based downstream interpretation of single-cell data.

### Application to Alzheimer’s disease single-nucleus RNA-seq dataset

To demonstrate its practical utility on real disease data, we applied **GatorSC** to a single-nucleus RNA-seq dataset from human entorhinal cortex (GEO: GSE138852) comprising 13,214 nuclei profiled from six individuals with AD and six age-matched controls [39]. In the learned latent space, **GatorSC** robustly segregates cells into five major brain cell populations, microglia, neurons, oligodendrocyte progenitor cells (OPCs), astrocytes, and oligodendrocytes (Figure 7A), in agreement with the original annotation of this dataset. The corresponding UMAP embedding shows that AD and control nuclei are well intermingled within each cell type, indicating that **GatorSC** captures biologically meaningful cell identities rather than trivially separating samples by disease status.

**Figure 7.**
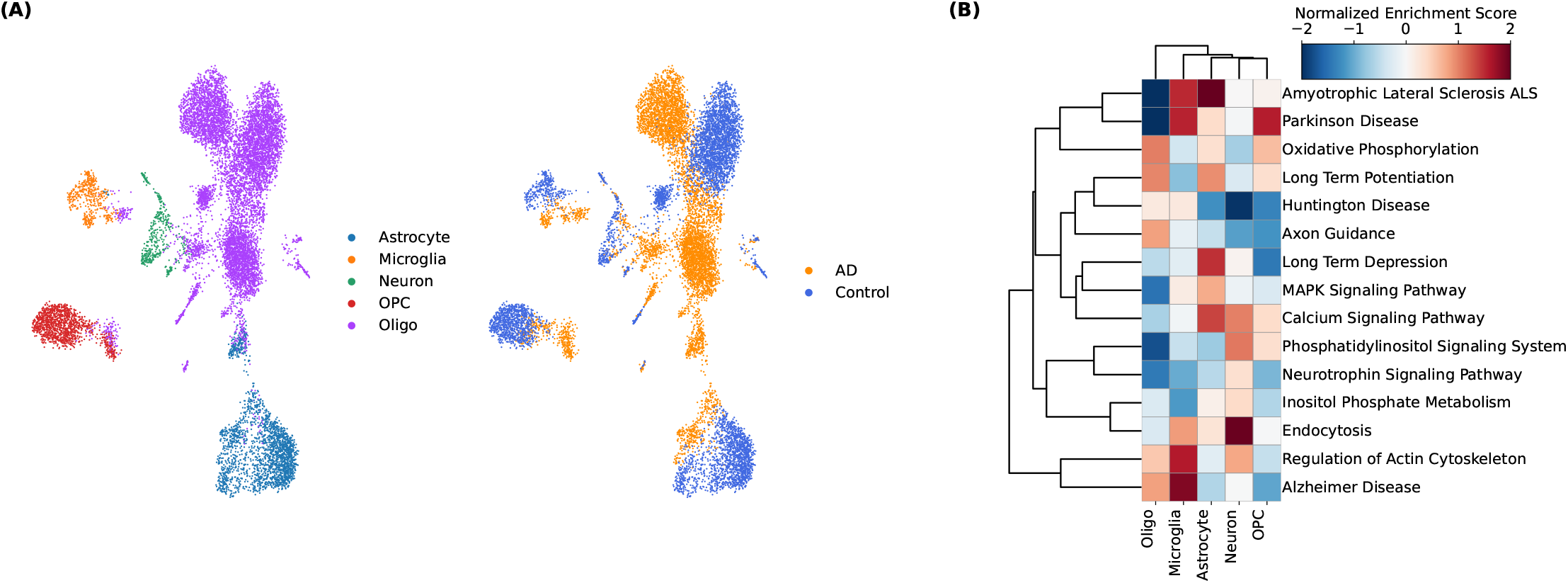
**Application of GatorSC to an Alzheimer’s disease single-nucleus RNA-seq dataset. (A) UMAP visualization of single-nucleus profiles, colored by GatorSC-inferred cell clusters (left) and by disease status (AD vs. control; right). (B) Heatmap of normalized enrichment scores for selected AD-related pathways in each major cell type, comparing AD versus control nuclei**.

Within each **GatorSC**-defined cell type, we then contrasted AD and control nuclei and performed pathway enrichment analysis (Figure 7B). Across all five cell populations, AD cells show strong positive enrichment for oxidative phosphorylation and for hallmark neurodegenerative disease pathways, including those associated with Alzheimer’s disease, Parkinson’s disease, Huntington’s disease, and amyotrophic lateral sclerosis. In addition, we observe pronounced cell-type-specific modulation of neuronal signaling pathways, such as axon guidance, long-term potentiation and depression, calcium signaling, and regulation of the actin cytoskeleton, indicating widespread perturbation of synaptic and neurodevelopmental programs in AD.

Notably, the MAPK (mitogen-activated protein kinase) signaling pathway shows a negative enrichment specifically in oligodendrocytes, whereas it is weakly perturbed or positively enriched in the other cell types (Figure 7B). This pattern suggests a relative suppression of MAPK-mediated regulation in myelinating oligodendrocytes compared to neurons, astrocytes, microglia, and OPCs, pointing to a distinct regulatory response of the oligodendrocyte lineage in AD. Together, these results demonstrate that **GatorSC** can be effectively applied to complex human snRNA-seq datasets to recover major brain cell classes and to dissect cell-type-specific pathway alterations and regulatory programs associated with Alzheimer’s disease.

## Conclusion

We introduced **GatorSC**, a unified representation learning framework for scRNA-seq that models multi-scale cell and gene structures with hierarchical graphs and fuses complementary views via a Mixture-of-Experts architecture. Trained with a dual self-supervised objective combining graph reconstruction and contrastive learning, **GatorSC** learns robust, biologically meaningful cell embeddings without task-specific supervision. Across 19 public datasets, **GatorSC** achieves consistently state-of-the-art performance on cell clustering, gene expression imputation, and cell-type annotation, and scales well to large, deeply annotated atlases while preserving fine-grained cellular structure. Beyond quantitative benchmarks, biological analyses (trajectory inference, marker-gene analysis, and a disease case study) further support the biological fidelity and interpretability of the learned embeddings, and ablation and sensitivity studies validate the contributions of hierarchical gene graphs, adaptive MoE fusion, and unified self-supervision.

## Data availability

The source code of **GatorSC** is available at https://github.com/zhangzh1328/GatorSC.

## Author contributions statement

Yuxi Liu and Zhenhao Zhang contributed equally to this work. Yuxi Liu: Conceptualization, Software, Formal analysis, Validation, Investigation, Methodology, Writing—original draft; Zhenhao Zhang: Data curation, Software, Formal analysis, Validation, Visualization, Methodology, Writing—original draft; Mufan Qiu: Software, Formal analysis, Validation; Song Wang: Writing—original draft; Flora D. Salim, Jun Shen, Tianlong Chen, Imran Razzak: Writing—review & editing; Fuyi Li: Conceptualization, Validation, Writing—review & editing; Jiang Bian: Conceptualization, Resources, Supervision, Funding acquisition, Project administration, Writing—review & editing.

**Key Points**

- GatorSC represents scRNA-seq with three hierarchical graphs to capture cell–cell and gene–gene structure at global and local scales.
- GatorSC uses a Mixture-of-Experts gating network to fuse multi-graph embeddings, trained with self-supervised reconstruction and contrastive learning.
- Across 19 public datasets, GatorSC delivers state-of-the-art clustering, imputation, and annotation, and supports trajectory, marker, and pathway analyses.

## Data pre-processing

**GatorSC** takes the scRNA-seq gene expression matrix as input. The scRNA-seq data are represented as a two-dimensional matrix, where rows correspond to cells, and columns to genes. Each cell is associated with an available label. The matrix is processed using the Scanpy toolkit. Preprocessing involves normalization and logarithmic transformation of the gene expression matrix.

## Implementation and parameter settings

GatorSC was implemented in Python (v3.9.12) with PyTorch (v1.11.2). For each scRNA-seq dataset, we randomly split cells into training/validation/test sets with a ratio of 80%/10%/10%. For hierarchical graph modeling, we used two attention heads and initialized the learnable thresholds as *ϕ*^(1)^ = 0.05, *ϕ*^(2)^ = 0.02, and *ϕ*^(3)^ = 0.03; the hop number was set to 2. For adaptive fusion of multi-level graph representations, the hidden dimension of both the GCN and MLP was set to 225, and the number of expert networks was set to 3. For unified self-supervision, the Bernoulli deletion and addition probabilities were both set to 0.4, and the maximum path length was set to 2. The temperature was set to *τ* = 0.25. The loss scaling coefficients were set to *α*_1_ = 0.55, *α*_2_ = 0.53, and *α*_3_ = 0.62; *β*_1_ = 0.90 and *β*_2_ = 0.85; *γ*_1_ = 0.76 and *γ*_2_ = 0.89; and *λ* = 0.68. We trained the model using the Adam optimizer with a learning rate of 10^−3^ and a batch size of 256. For clustering, we applied K-means (scikit-learn default settings) to the learned embeddings, setting the number of clusters to the ground-truth number of cell types for each dataset. For other clustering baselines, we used the default hyperparameters provided by the corresponding packages. All hyperparameters were selected via grid search based on validation performance. All experiments were conducted on a workstation equipped with an NVIDIA RTX 4090 GPU (24 GB memory).

## Evaluation metrics

For cell clustering performance, we utilize adjusted Rand index (ARI), normalized mutual information (NMI), clustering accuracy (ACC), and homogeneity score (Homogeneity) as evaluation metrics.

ARI measures the agreement between the predicted and reference clusterings while accounting for randomness, and is therefore particularly useful when the class distributions are unbalanced or when clusters overlap. The ARI between the ground truth **Y** and the predicted **Ŷ** can be written as follows:

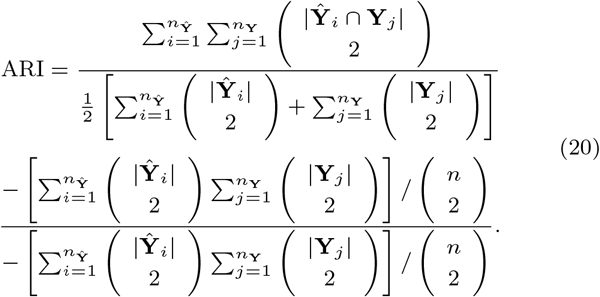

where *n*_**Y**_ and *n*_**Ŷ**_ denote the numbers of clusters in **Y** and **Ŷ**, respectively, and *n* is the total number of cells.

NMI quantifies the mutual dependence between the predicted and ground truth labels, while the normalization alleviates the influence of imbalanced label distributions. The NMI between **Y** and **Ŷ** be written as follows:

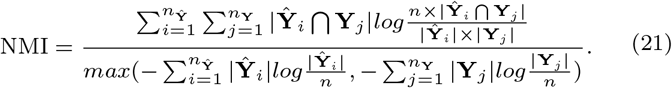

ACC measures the proportion of correctly assigned cells after optimally matching predicted clusters to ground-truth labels. The ACC between **Y** and **Ŷ** can be written as follows:

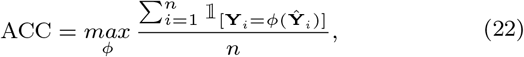

where *ϕ* is a mapping from predicted cluster indices to ground-truth labels, and 𝟙_[·]_ denotes the indicator function.

Finally, the homogeneity score evaluates the extent to which each cluster contains only cells from a single ground-truth class as follows:

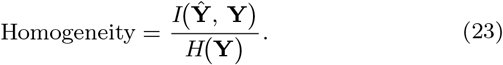

where *H*(**Y**) is the entropy of **Y** and *I*(**Ŷ**, **Y**) is the mutual information between **Ŷ** and **Y**. The score ranges from 0 to 1, with 1 indicating perfectly pure clusters (each cluster containing cells from only one class), whereas lower values indicate mixed clusters.

For the gene expression imputation, we adopt widely used metrics: Pearson correlation coefficient (PCC) and L1 loss.

Specifically, PCC measures the linear correlation between the imputed values and ground truth expression matrix, with higher scores suggesting better preservation of global structural patterns in the data. The PCC between the ground truth **X** and the imputed value 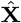 can be written as follows:

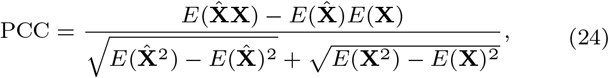

where *E*(·) is the expected value. The L1 loss captures the average absolute deviation between the predicted and true values, providing a robust measure of local imputation accuracy. The L1 between **X** and 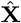 can be written as follows:

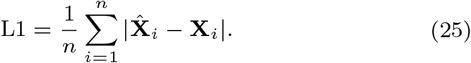

For the cell-type annotation, we report the F1 score (F1), classification accuracy (ACC), and Matthews correlation coefficient (MCC), three widely used metrics for evaluating classification performance. Given the numbers of true positives (TP), false positives (FP), false negatives (FN), and true negatives (TN), these metrics can be written as:

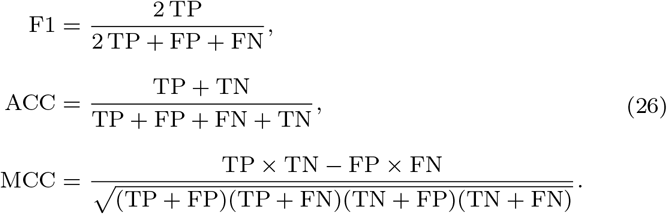

## Cell-type annotation across annotation levels and dataset scales

To further examine the robustness of **GatorSC** for cell-type annotation, we considered datasets that vary in both annotation depth and dataset size. Specifically, we used the Allen Mouse Brain (AMB) dataset with three nested annotation levels containing 3, 16, and 92 cell populations (AMB3, AMB16, and AMB92) to assess performance as labels become increasingly fine-grained. We also included Tabula Muris (TM) and Zheng 68K, two large-scale scRNA-seq datasets with *>* 50,000 cells, to evaluate scalability in high-throughput scenarios. As shown in Figure 8, **GatorSC** achieves state-of-the-art or comparable F1, ACC, and MCC scores across all three AMB levels. While all methods drop when moving from AMB3 to the most challenging AMB92 setting, **GatorSC** retains a clear advantage or remains among the top performers, suggesting that its representations preserve meaningful structure under fine-grained label schemes. On TM and Zheng 68K, **GatorSC** again matches or outperforms existing classifiers across all metrics, indicating that the hierarchical graph representation and Mixture-of-Experts fusion support accurate annotation of subtle subpopulations and scale effectively to large datasets. Overall, these results demonstrate that **GatorSC** provides a consistent and robust annotation framework across a wide range of annotation resolutions and dataset sizes.

**Figure 8.**
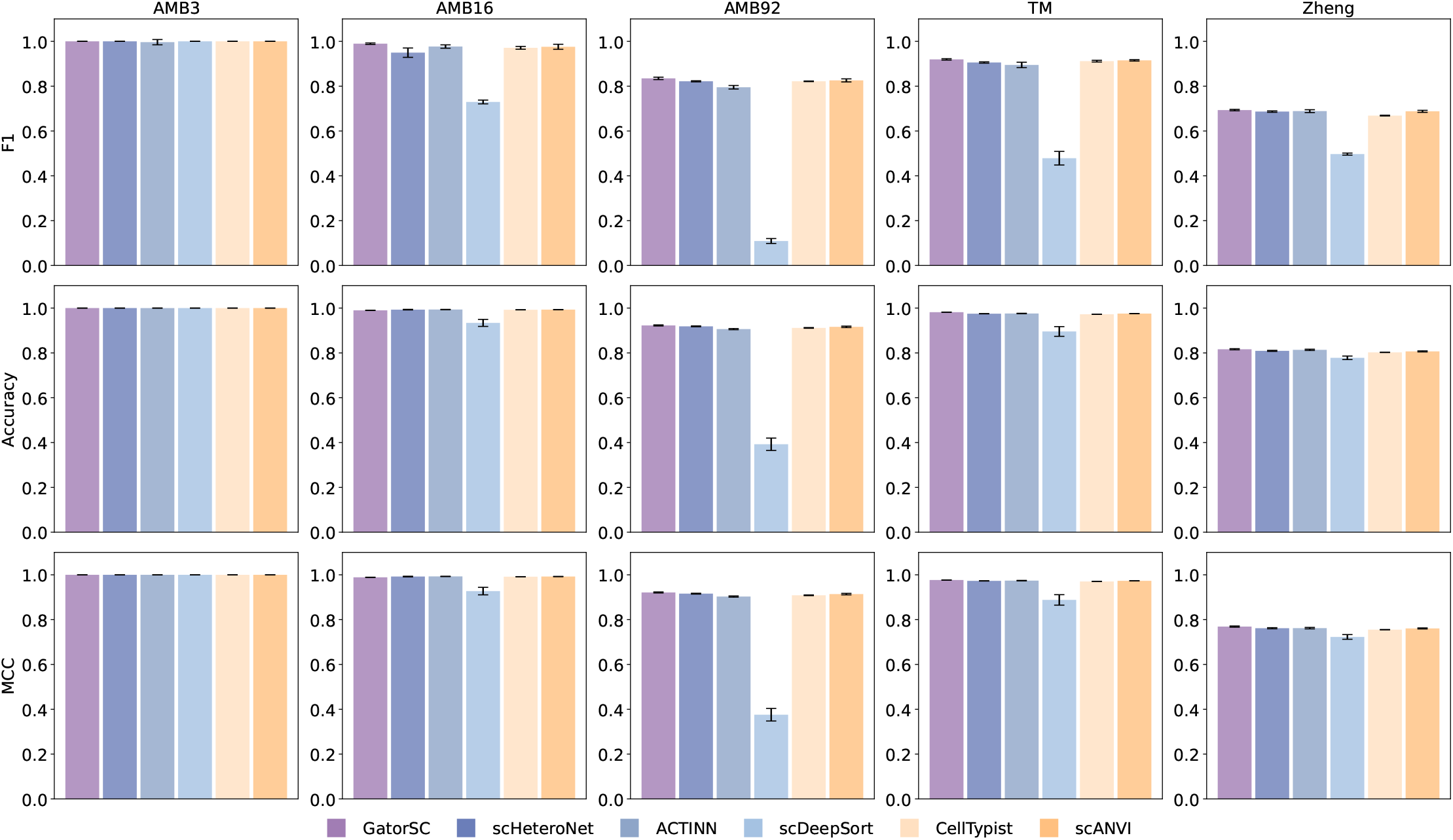
**Cell-type annotation performance of GatorSC and competing methods on five scRNA-seq datasets with varying annotation granularity and dataset size. Columns correspond to the three annotation levels of the Allen Mouse Brain dataset (AMB3, AMB16, AMB92; with 3, 16, and 92 cell populations, respectively), the Tabula Muris (TM) dataset, and the Zheng 68K PBMC dataset. Bars show F1 score (top row), overall accuracy (middle row), and Matthews correlation coefficient (MCC, bottom row) for each method. Across most datasets and metrics, GatorSC achieves state-of-the-art or comparable performance, and higher values of all three metrics indicate better cell-type annotation**.

### t-SNE Visualization of Learned Latent Representations for GatorSC and Baseline Methods

Figure 9 presents the t-SNE visualization of the latent embeddings learned by all baseline methods and **GatorSC** on the Emont and Quake-Trachea datasets. For each method, we first extract the intermediate low-dimensional representation of each cell and then apply t-SNE to obtain a two-dimensional embedding, on which cells are colored according to their ground-truth cell-type labels. This allows a direct qualitative comparison of how well different models separate cell types in the latent space. Across both datasets, the clusters generated by **GatorSC** are more compact and clearly separated than those of competing methods. On Emont, **GatorSC** yields distinct islands for most cell types and achieves the highest ARI (0.763), outperforming strong baselines such as scDSC (ARI = 0.715) and scSimGCL (ARI = 0.702). On the Quake-Trachea dataset, **GatorSC** further improves the agreement with ground truth with an ARI of 0.801, again exceeding scSimGCL (ARI = 0.778) and scGPCL (ARI = 0.745). In contrast, several baselines show noticeable mixing between cell types in the t-SNE space. Methods such as Desc and scDeepCluster form partially entangled clusters, with blurred boundaries between neighboring populations on both datasets, consistent with their moderate ARI values. Latent-variable models including scVI and DeepBID separate some major populations but still show overlap in regions where **GatorSC** produces well-isolated clusters. The graph-based approach graph-sc shows limited separation on Emont, where most cells collapse into a single diffuse cloud, consistent with its lower ARI, and only partially recovers the cluster structure on Quake-Trachea. Even the stronger baselines scDSC, scGPCL, and scSimGCL, while yielding more discernible groups, tend to produce clusters that are less compact and more fragmented than those of **GatorSC**. Taken together, these t-SNE visualizations indicate that **GatorSC** learns a latent space in which biologically meaningful cell populations are more cleanly separated than with existing methods.

**Figure 9.**
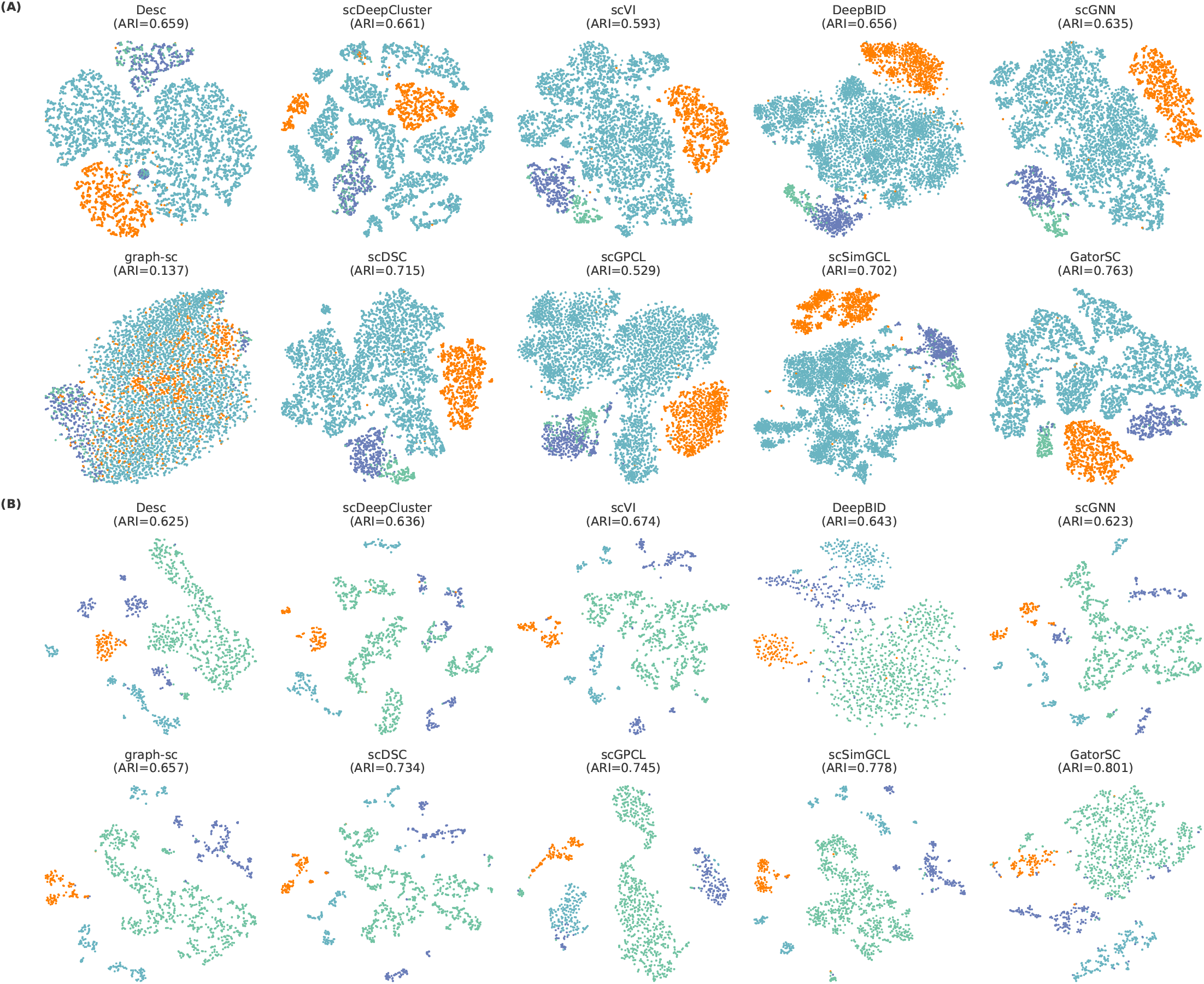
**Visualization analysis results of baselines and GatorSC on the (A) Emont and (B) Quake-Trachea datasets**.

## Ablation Study

To quantify the contribution of each component in **GatorSC**, we conducted an ablation study comparing the full model with six variants, as shown in Figure 10. In **GatorSC**_*α*_, we removed the MoE fusion and instead concatenated *Z*^(1)^, *Z*^(2)^, and *Z*^(3)^, followed by a linear projection back to the original latent space. **GatorSC**_*β*_ and **GatorSC**_*γ*_ retained only the contrastive or reconstruction objective, respectively, while discarding the complementary self-supervised loss terms. **GatorSC**_*δ*_ removed the local gene–gene graph and fused only *Z*^(1)^ and *Z*^(2)^, whereas **GatorSC**_*ε*_ removed the global gene–gene graph and fused *Z*^(1)^ and *Z*^(3)^. Finally, **GatorSC**_*ζ*_ used only the global cell–cell graph, directly treating *Z*^(1)^ as the final representation. Across 14 benchmark scRNA-seq datasets, the full **GatorSC** consistently achieves the best ARI, NMI, ACC, and homogeneity scores, whereas all ablated variants exhibit noticeable performance degradation. The performance gap between **GatorSC** and **GatorSC**_*α*_ indicates that MoE-based adaptive fusion is more effective than a simple concatenation-plus-linear layer, highlighting the benefit of data-dependent expert weighting. The inferior results of **GatorSC**_*β*_ and **GatorSC**_*γ*_ demonstrate that contrastive and reconstruction objectives provide complementary supervisory signals, and that removing either objective harms clustering quality. Moreover, the performance drops observed for **GatorSC**_*δ*_ and **GatorSC**_*ε*_ confirm that both global and local gene–gene graphs contribute non-redundant information beyond the cell–cell graph. Finally, **GatorSC**_*ζ*_, which relies solely on the global cell–cell graph, yields suboptimal performance, highlighting the importance of jointly modeling hierarchical gene–gene structure and employing dual self-supervised learning to obtain robust and discriminative cell representations.

**Figure 10.**
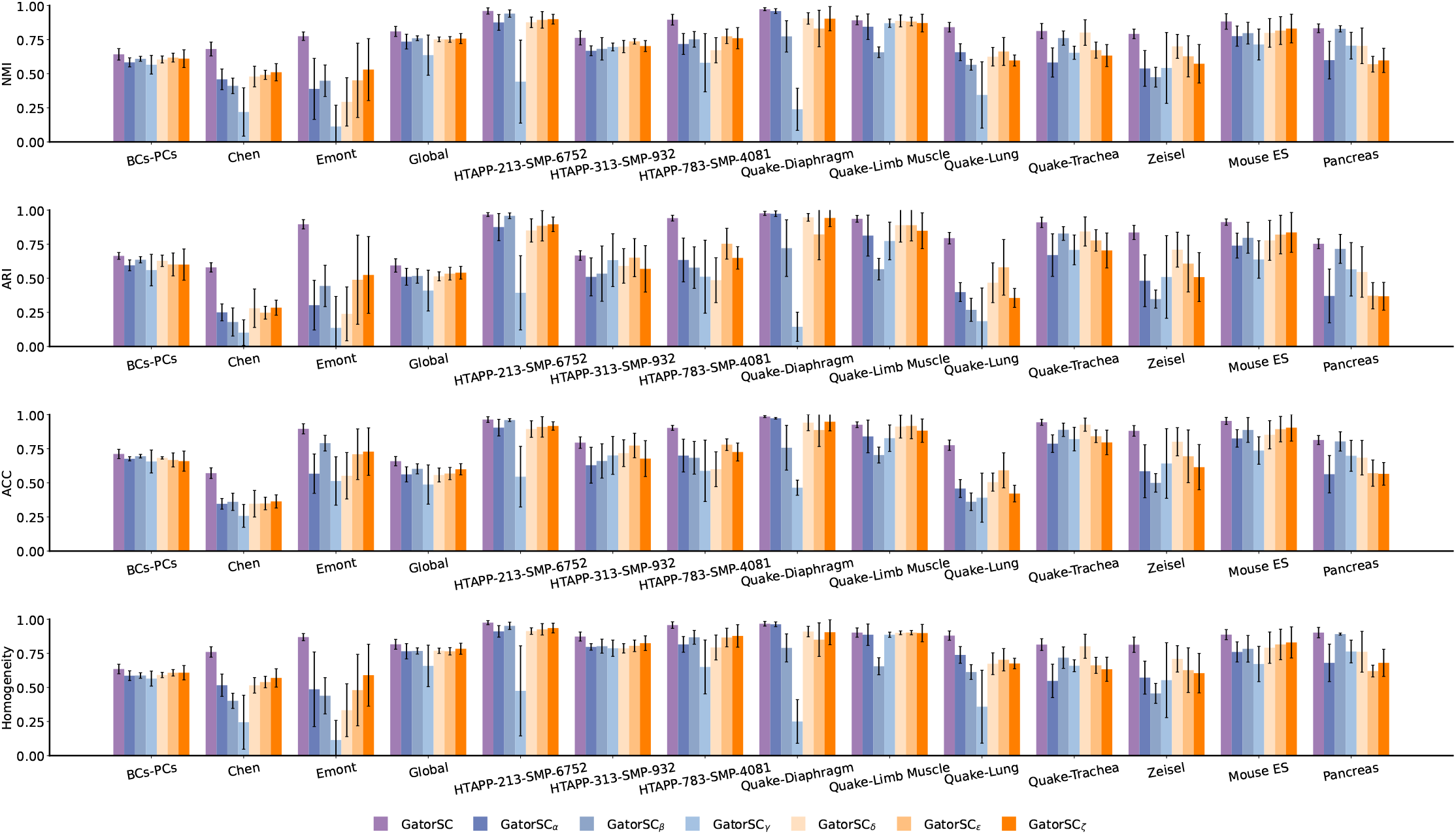
**Clustering performance comparison across 14 scRNA-seq datasets between GatorSC and its six variants**.

## Hyperparameter sensitivity analysis

We systematically evaluated the sensitivity of **GatorSC** to its key hyperparameters, namely the *k*-hop radius *n*_*k*_, the path-wise neighborhood size *n*_*p*_, the contrastive temperature *τ*, and the dropout rate *p*_drop_. For each hyperparameter, we varied its value over a wide range and assessed the resulting clustering quality using ARI, NMI, ACC, and homogeneity across the 14 benchmark scRNA-seq datasets. As summarized in Figure 11, the performance curves remain largely flat over broad intervals for all four hyperparameters, indicating that **GatorSC** maintains stable clustering accuracy under substantial variation in its configuration. While very extreme settings can lead to a mild decrease in performance on a subset of datasets, we do not observe any abrupt performance collapse or catastrophic degradation. Instead, comparable performance is consistently achieved across a continuum of reasonable choices for *n*_*k*_, *n*_*p*_, *τ*, and *p*_drop_, and the qualitative trends are aligned across all evaluation metrics. These results demonstrate that **GatorSC** is robust to hyperparameter variation and does not rely on carefully tuned, dataset-specific configurations, which reduces the burden of hyperparameter selection and facilitates its deployment to new scRNA-seq datasets.

**Figure 11.**
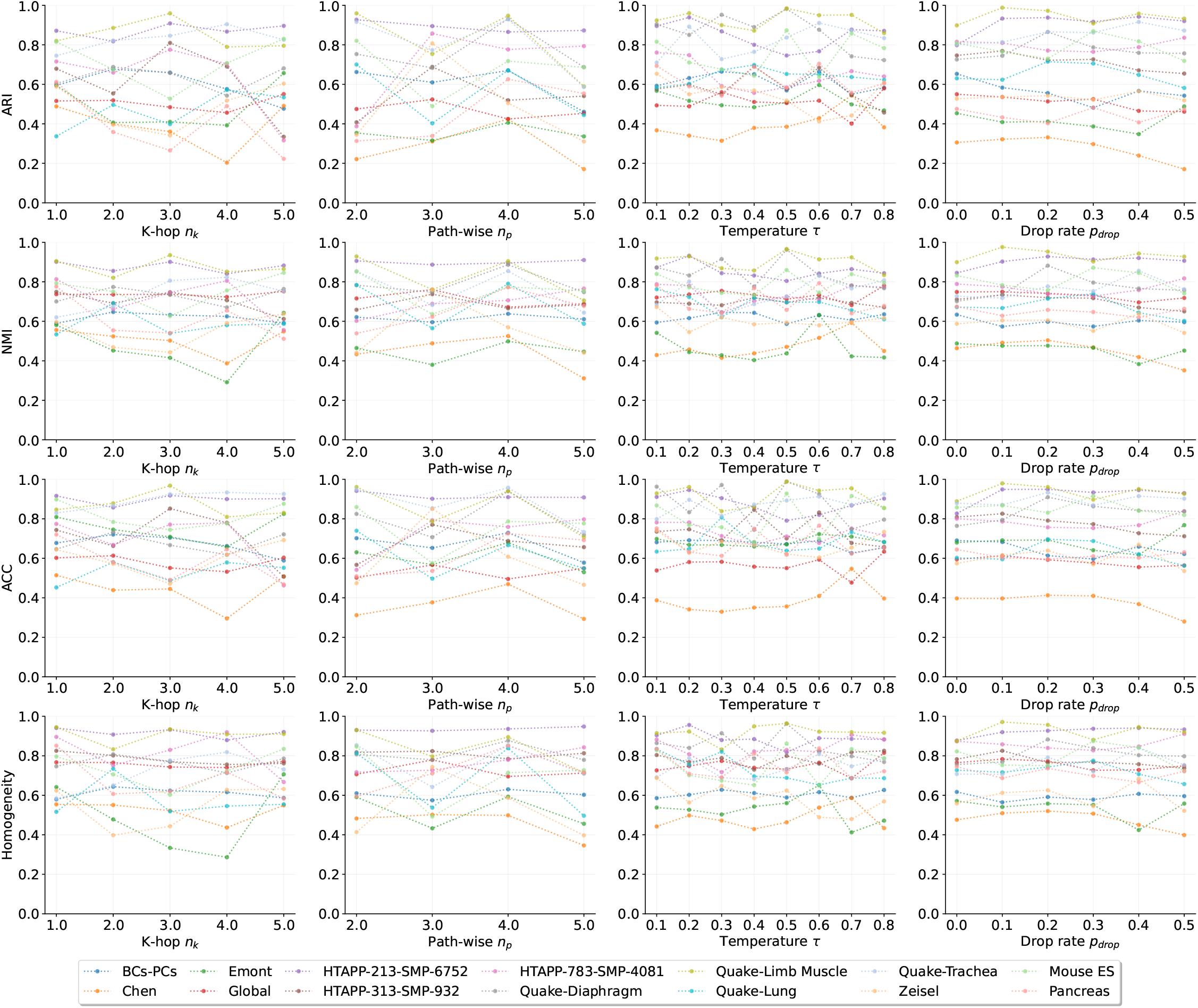
**Hyperparameter analysis results using four clustering evaluation metrics: ARI, NMI, ACC, and Homogeneity on 14 scRNA-seq datasets**.

